# Modeling the temporal dynamics of the gut microbial community in adults and infants

**DOI:** 10.1101/212993

**Authors:** Liat Shenhav, Ori Furman, Leah Briscoe, Michael Thompson, Itzhak Mizrahi, Eran Halperin

## Abstract

Given the highly dynamic and complex nature of the human gut microbial community, the ability to identify and predict time-dependent compositional patterns of microbes is crucial to our understanding of the structure and function of this ecosystem. One factor that could affect such time-dependent patterns is microbial interactions, wherein community composition at a given time point affects the microbial composition at a later time point. However, the field has not yet settled on the degree of this effect. Specifically, it has been recently suggested that only a minority of the operational taxonomic units (OTUs) depend on the microbial composition in earlier times. To address the issue of identifying and predicting temporal microbial patterns we developed a new model, *MTV-LMM* (Microbial Temporal Variability Linear Mixed Model), a linear mixed model for the prediction of the microbial community temporal dynamics based on the community composition at previous time stamps. *MTV-LMM* can identify time-dependent microbes in time series datasets, which can then be used to analyze the trajectory of the microbiome over time. We evaluated the performance of *MTV-LMM* on three human microbiome time series datasets, and found that *MTV-LMM* significantly outperforms all existing methods for microbiome time series modeling. Particularly, we demonstrate that the effect of the microbial composition in previous time points on the abundance levels of an OTU at a later time point is underestimated by a factor of at least 10 when applying previous approaches. Using *MTV-LMM*, we demonstrate that a considerable proportion of the human gut microbiome, both in infants and adults, has a significant time-dependent component that can be predicted based on microbiome composition in earlier time points. This suggests that microbiome composition at a given time point is a major factor in defining future microbiome composition and that this phenomenon is considerably more common than previously reported for the human gut microbiome.

## Introduction

There is increasing recognition that the human gut microbiome is a contributor to many aspects of human physiology and health including obesity, non-alcoholic fatty liver disease, inflammatory diseases, cancer, metabolic diseases, aging, and neurodegenerative disorders [1–14]. This suggests that the human gut microbiome may play important roles in the diagnosis, treatment, and ultimately prevention of human disease. These applications require an understanding of the temporal variability of the microbiota over the lifespan of an individual particularly since we now recognize that our microbiota is highly dynamic, and that the mechanisms underlying these changes are linked to ecological resilience and host health [15–17].

Due to the lack of data and insufficient methodology, we currently have major gaps in our understanding of fundamental mechanisms related to the temporal behavior of the microbiome. Critically, we currently do not have a clear characterization of how and why our gut microbiome varies in time, and whether these dynamics are consistent across humans. It is also unclear whether we can define’stable’ or ‘healthy’ dynamics as opposed to ‘abnormal’ or ‘unhealthy’ dynamics, which could potentially reflect an underlying health condition or an environmental factor affecting the individual, such as antibiotics exposure or diet. Moreover, there is no consensus as to whether the gut microbial community structure varies continuously or jumps between discrete community states, and whether or not these states are shared across individuals [18, 19]. Notably, recent work [20] suggests that the human gut microbiome composition is dominated by environmental factors rather than by host genetics, emphasizing the dynamic nature of this ecosystem.

The need for understanding the temporal dynamics of the microbiome and its interaction with host attributes have led to a rise in longitudinal studies that record the temporal variation of microbial communities in a wide range of environments, including the human gut microbiome. These time series studies are enabling increasingly comprehensive analyses of how the microbiome changes over time, which are in turn beginning to provide insights into fundamental questions about microbiome dynamics [16, 17, 21].

One of the most fundamental questions that still remains unanswered is to what degree the microbial community in the gut is deterministically dependent on its initial composition. More generally, it is unknown to what degree the microbial composition of the gut at a given time determines the microbial composition at a later time. Additionally, there is only preliminary evidence of the long-term effects of early life events on the gut microbial community composition, and it is currently unclear whether these long-term effects traverse through a predefined set of potential trajectories [21, 22].

In order to answer these questions, it is important to quantify the dependency of the microbial community at a given time on past community composition [23,24]. This task has been previously studied in theoretical settings. Specifically, the generalized Lotka-Volterra family of models predict changes in community composition through defined species-species or species-resource interaction terms, and are popular for describing internal ecological dynamics such as external factors (e.g., diet and host behavior [17]), species-species interactions (e.g., cross-feeding or successional turnover), and host-species interactions (e.g., immune system regulation or host physiology) [25–27]. The generalized Lotka-Volterra models are deterministic and fairly straightforward to interpret, but little is known about the relative importance of purely autoregressive factors (a stochastic process in which future values are a function of the weighted sum of past values) in driving gut microbial dynamics. Moreover, it has been recently shown that even microbial communities that are in an equilibrium state do not correspond to the generalized Lotka-Volterra model [28], which emphasizes the need for new models that provide insights about the microbial temporal dynamics.

Using their sparse vector autoregression (sVAR) model, Gibbons et al. [24] have recently found evidence that the human gut microbial community has two dynamic regimes: autoregressive and non-autoregressive. The autoregressive regime includes operational taxonomic units (OTUs) that are affected by the community composition at previous time points, while the non-autoregressive regime includes OTUs that are completely stochastic, i.e. their appearance in a specific time is random and does not depend on the past, and these OTUs in particular will not behave according to models such as the generalized Lotka-Volterra. Additionally, Ridenhour et al. [29] suggest modeling the OTU read counts along time using Poisson regression (ARIMA Poisson) in order to infer the temporal interactions within the microbial community. sVAR and ARIMA Poisson assume linear and log-linear dynamics, respectively, and are built around an autoregressive model that uses the last p timestamps (also known as AR(p)). In contrast, the generalized Lotka-Volterra family of models assumes that the differenced log-transformed data is AR(1). The relation between AR(p) and AR(1) in this scenario has been demonstrated in Gibbons et al., 2017 [24].

In this paper, we show that previous studies substantially underestimate the autoregressive component of the gut microbiome. In order to quantify the dependency of an OTU on the past composition of the microbial community, we developed a method, Microbial community Temporal Variability Linear Mixed Model (*MTV-LMM*). Our method is based on a linear mixed model, a heavily used tool in statistical genetics and other areas of genomics [30, 31]. We define the term time-explainability, which quantifies the overall association between the microbial relative abundance in the present time and microbial composition in the past after accounting for the effect of the individual host. Put differently, for each OTU, we quantify the temporal effect - the effect of the community composition in previous time points on the OTU abundance in the current time point.

Similarly to previous approaches, e.g. sVAR, *MTV-LMM* also assumes linear dynamics and is built around an AR(p) type of model. Thus, *MTV-LMM* is related to previous biological models, i.e. the generalized Lotka-Volterra, in much the same way as previous works such as sVAR and ARIMA Poisson (see Supplementary for a characterization of the relation between *MTV-LMM* and the generalized Lotka-Volterra family of models). Additionally, *MTV-LMM* has a natural interpretation: it assumes that the abundance of a specific OTU at time *t* + 1 is affected by the abundance levels of many OTUs at previous time points. The underlying assumption is that each of these effects is very small, and we thus do not attempt to infer it but rather integrate over all possible values of these effects. This integration induces regularization and hence reduces the number of parameters estimated by the model considerably.

Our approach has a few notable advantages. First, unlike the sVAR model our approach models all the individual hosts simultaneously, thus leveraging the information across an entire population while adjusting for the individual’s effect, e.g. the host’s genetics or environment. This provides *MTV-LMM* an increased power to detect temporal dependencies, as well as the ability to quantify the consistency of dynamics across individuals. The Poisson regression method suggested by Ridenhour et al. [29] also utilizes the information from all individuals, but does not account for the individual effects, which may result in an inflated autoregressive component. Second, *MTV-LMM* can serve both as a feature selection method (selecting only the OTUs affected by time) and as a prediction model. The ability to identify the time-dependent OTUs is crucial when fitting a time series model to study the microbial community temporal dynamics. Finally, we demonstrate that *MTV-LMM* can serve as a standalone prediction model that outperforms existing prediction methods by an order of magnitude in predicting the OTU abundance.

We applied *MTV-LMM* to simulated data as well as to three real time series datasets of the gut microbiome (David et al. [17], Caporaso et al. [16], and DIABIMMUNE [21]). These datasets contain longitudinal abundance OTU data (using 16S rRNA gene sequencing). Notably, *MTV-LMM* is agnostic to the input data type used to represent species abundance. Our results show that, compared to previous approaches, *MTV-LMM* has a substantially improved prediction accuracy of the OTU relative abundance levels at the next timestamp, indicating that the underlying model used by *MTV-LMM* better predicts the future microbiome composition. Moreover, we show that, on average, the time-explainability (the variance of the abundance level explained by previous timestamps) of an OTU is an order of magnitude larger than previously estimated for these datasets. This can provide insights for future extensions of temporal ecological models such as the generalized Lotka-Volterra family of models [32].

Finally, with *MTV-LMM* one can address the formative question surrounding the proportion of microbes that depend on past composition of the microbial community. Unlike previously thought, we found that a considerable proportion of microbes in the human gut depend on early microbiome composition and that the degree of this dependency varies between microbes – ranging from complete to moderate dependence.

## Results

### A brief description of *MTV-LMM*

We begin with an informal description of the main idea and utility of *MTV-LMM*, but a more comprehensive description can be found in the Methods. *MTV-LMM* is motivated by our assumption that the temporal changes in abundance levels of a specific OTU *j* are a time-homogeneous high-order Markov process. *MTV-LMM* models the transitions of this Markov process by fitting a sequential linear mixed model (LMM) to predict the relative abundance level of an OTU at a given time point, given the microbial community composition at previous time points. Intuitively, the linear mixed model correlates the similarity between the microbial community composition across different timestamps with the similarity of the OTU abundance levels at the next time points. The input to *MTV-LMM* is the microbial community abundance at time points *{*1*,…,t −* 1*}*, and its output is the prediction of the relative abundance, for each OTU, at time *t*. In order to apply linear mixed models, *MTV-LMM* generates a *temporal kinship matrix*, which represents the similarity between every pair of samples across time, where a sample is a normalization of the abundances of the OTUs at a given time point for a given individual (see Methods). When predicting the abundance level of a specific OTU *j* at time *t*, the model uses both the global state of the entire microbial community in the last *q* time points, as well as the relative abundance levels of OTU *j* in the previous *p* time points. The parameters *p* and *q* are determined by the user, or they can be determined using a cross-validation approach; a more formal description of their role is provided in the Methods.

*MTV-LMM* has the advantage of increased power due to a low number of parameters coupled with an inherent regularization mechanism, similar in essence to the widely used ridge regularization, which provides a natural interpretation of the model (see Methods). Specifically *MTV-LMM* has six parameters: the two temporal parameters (*p* and *q*), coefficients of the fixed effects, and variance of the random effects, individual effects, and environmental effects (see Methods).

### Model Evaluation

We evaluated *MTV-LMM* by testing its accuracy in predicting the abundance levels of all the OTUs at a future time point using real data. Such evaluation is guaranteed to avoid overfitting, since the future data points are held out from the algorithm. In addition to the prediction accuracy on real time series data, we also validated *MTV-LMM* using simulations. We constructed artificial microbial community data using a randomized interaction network and calculated the accuracy in predicting the abundance level of all the OTUs at a future time (see Supplementary).

We used real time series data from three different datasets, each composed of longitudinal abundance OTU data. These three datasets are David et al. (2 adult donors - DA, DB - average 250 time points per individual), Caporaso et al. (2 adult donors - M3, F4 - average 231 time points per individual), and the DIABIMMUNE dataset (39 infant donors - average 28 time points per individual). In these datasets, *p* and *q* were estimated using a training set, and ranged between 0 and 3 per OTU. See Methods for further details and justification for the use of multiple time points in the model (*p* and *q ≥* 1).

We compared the results of *MTV-LMM* to the approaches that have been used in the past for temporal microbiome modeling, namely the AR(1) model (see Methods), the sparse vector autoregression model sVAR [24] and the ARIMA (1, 0, 0) Poisson regression [29]. Overall, *MTV-LMM*’s prediction accuracy is higher than AR’s (Supplementary Table S1) and significantly outperforms both the sVAR method and the Poisson regression method across all datasets, using real and simulated time-series data (Fig. 1 and Supplementary Fig S1, S2). Therefore, we conclude that *MTV-LMM* is more suitable for estimating the autoregressive component in longitudinal microbial data.

**Figure 1.**
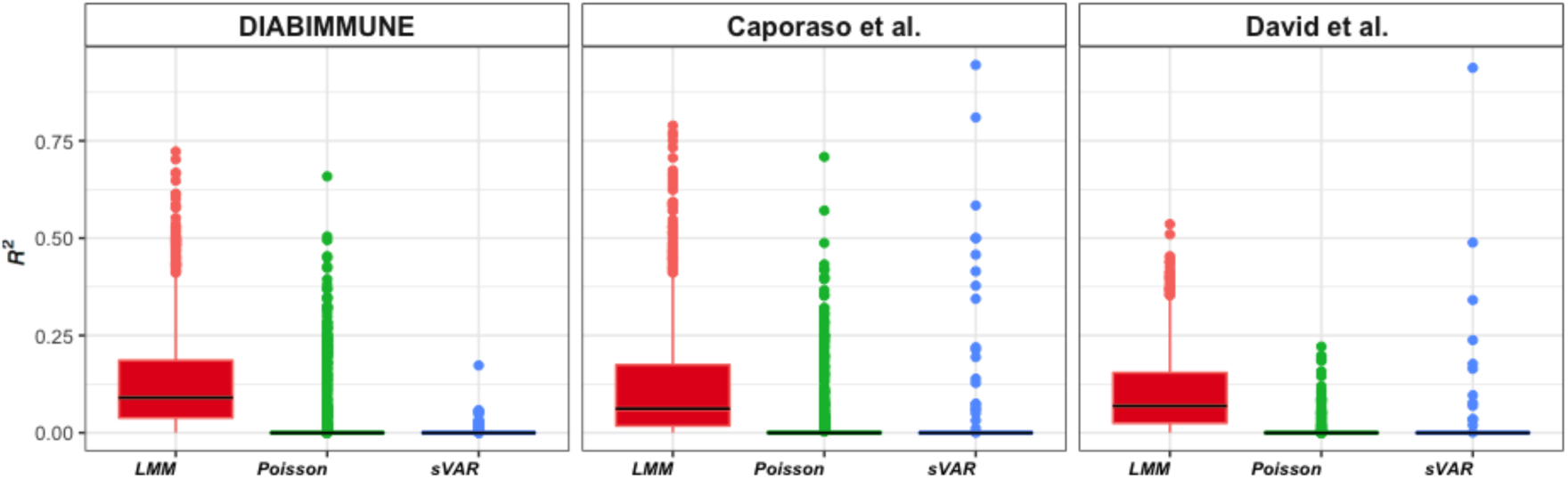
*MTV-LMM* outperforms all other methods in prediction accuracy (*R*^2^) and autoregressive detection. The *MTV-LMM* predictions are in red, sVAR in green, and ARIMA Poisson regression in blue.

For over 96% of the OTUs, the fraction of variance explained by previous composition of the microbial community (calculated by the squared correlation between the true and predicted abundance), estimated by *MTV-LMM* is significantly greater than that reported previously by sVAR [24] and ARIMA Poisson regression [29]. *MTV-LMM* explained on average 12.8% of the variance in the DIABIMMUNE dataset, 12.3% of the variance in Caporaso et al. (2011) and 10.4% in David el al.

In contrast, sVAR and ARIMA Poisson regression explained on average only 1.783% and 1.95% of the variance in the DIABIMMUNE data set respectively, 0.58% and 1.5% of the variance in Caporaso et al. (2011), and 0.28% and 0.4% of the variance in David el al.

### Inference on the estimated interaction matrix

We applied *MTV-LMM* on the DIABIMMUNE infant dataset and estimated the species-species interactions matrix across all individuals, using 1440 OTUs that passed a preliminary screening according to temporal presence-absence patterns (see Methods). We found that most of these effects are close to zero, implying a sparse interaction pattern. Next, we applied a principal component analysis (PCA) on the estimated species-species interactions and found a strong phylogenetic structure (PerMANOVA P-value = 0.001) suggesting that closely related species have similar interaction patterns within the microbial community (Fig. 2). These findings are supported by Thompson et al. [33], who suggest that ecological interactions are phylogenetically conserved, where closely related species interact with similar partners. Gomez et al. [34] tested these assumptions on a wide variety of hosts and found that generalized interactions can be evolutionary conserved.

**Figure 2.**
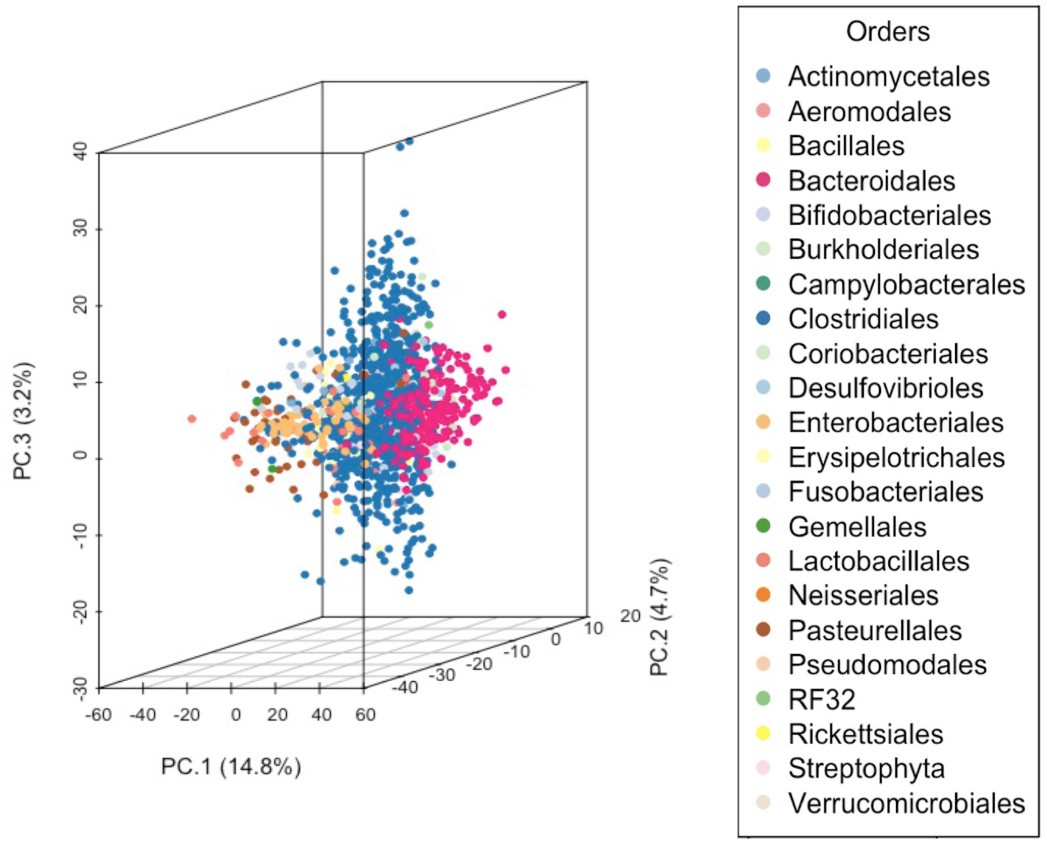
Phylogenetic structure in the species-species interactions matrix estimated by *MTV-LMM* on the DIABIMMUNE dataset, uncovered by principal component analysis, suggesting that closely related species may have similar interaction patterns within the microbial community. Shown on the axes are the percentage of variance explained by each principal component.

We note that the interactions matrix estimated by *MTV-LMM* should be interpreted with caution since the number of possible interactions is quadratic in the number of species, and it is therefore infeasible to infer with high accuracy all the interactions. However, we can still aggregate information across species or higher taxonomic levels to uncover global patterns of the microbial composition dynamics (e.g., principal component analysis).

### Time-explainability as a measure of the autoregressive component in the microbial community

In order to address the fundamental question regarding the gut microbiota temporal variation, we quantify its autoregressive component. Namely, we quantify to what degree the relative abundance of different OTUs can be inferred based on the composition at previous time points. In statistical genetics, the fraction of phenotypic variance explained by genetic factors is called heritability and is typically evaluated under an LMM framework [30]. Intuitively, linear mixed models estimate heritability by measuring the correlation between the genetic similarity and the phenotypic similarity of pairs of individuals. We used *MTV-LMM* to define an analogous concept that we term *time-explainability*, which corresponds to the fraction of OTU variance explained by the microbiome composition in previous time points. Specifically, time-explainability in linear mixed models measures the correlation between the similarity of the microbial community composition, across different timestamps and individuals, and the similarity of the OTU abundance levels at the next time points (see Methods for details).

We next estimated the time-explainability of the OTUs in each data set, using the parameters *q* = 1*,p* = 0. The resulting model, including only the effect of the microbial community, corresponds to the formula: *OTU*_*t*_ = microbiome community_(*t−*1)_ + individual effect_(*t−*1)_ + unknown effects. Notably, time-explainability can be estimated fairly accurately, with an average 95% exact confidence interval width of 23.7%, across all different time-explainability levels and across all datasets (Methods).

Of the OTUs we examined, we identified a large portion of them to have a statistically significant time-explainability component across datasets. Specifically, we found that over 85% of the OTUs included in the temporal kinship matrix are significantly explained by the time-explainability component, with estimated time-explainability average levels of 23% in the DIABIMMUNE infant dataset (sd = 15%), 21% in the Caporaso et al. (2011) dataset (sd = 15%) and 14% in the David el al. dataset (sd = 10%) (Fig. 3, Supplementary Fig. S3). Notably, the time-explainability estimates across datasets are highly correlated to the fraction of variance explained by the predictions *R*^2^ (Supplementary Fig. S4).

**Figure 3.**
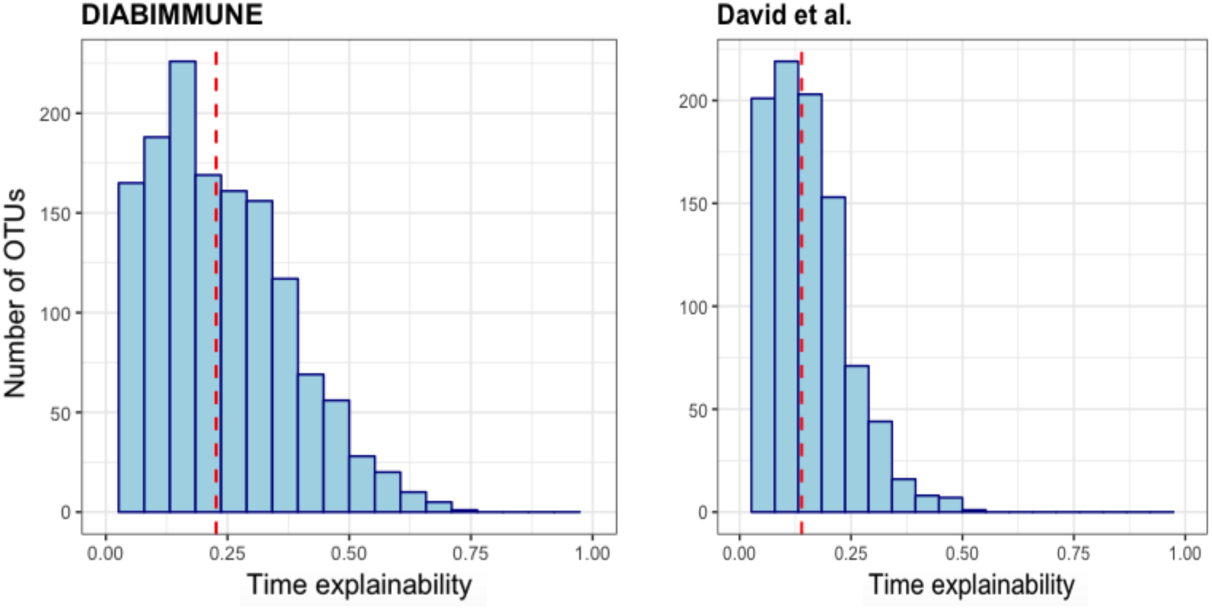
Time-explainability distribution in the DIABIMMUNE infant dataset (left) and David et al. adult dataset (right). The average time-explainability (denoted by a dashed line) in the DIABIMMUNE cohort is 23% and in David et al. is 14%.

### Non-autoregressive dynamics contain phylogenetic structure

As a secondary analysis, we aggregated the *MTV-LMM* time-explainability by taxonomic order, and found that in some orders (*non-autoregressive orders*) all OTUs are non-autoregressive, while in other orders (*mixed orders*) we observe the presence of both autoregressive and non-autoregressive OTUs (Fig. 4), where an autoregressive OTU is defined as an OTU having a statistically significant time-explainability component.

**Figure 4.**
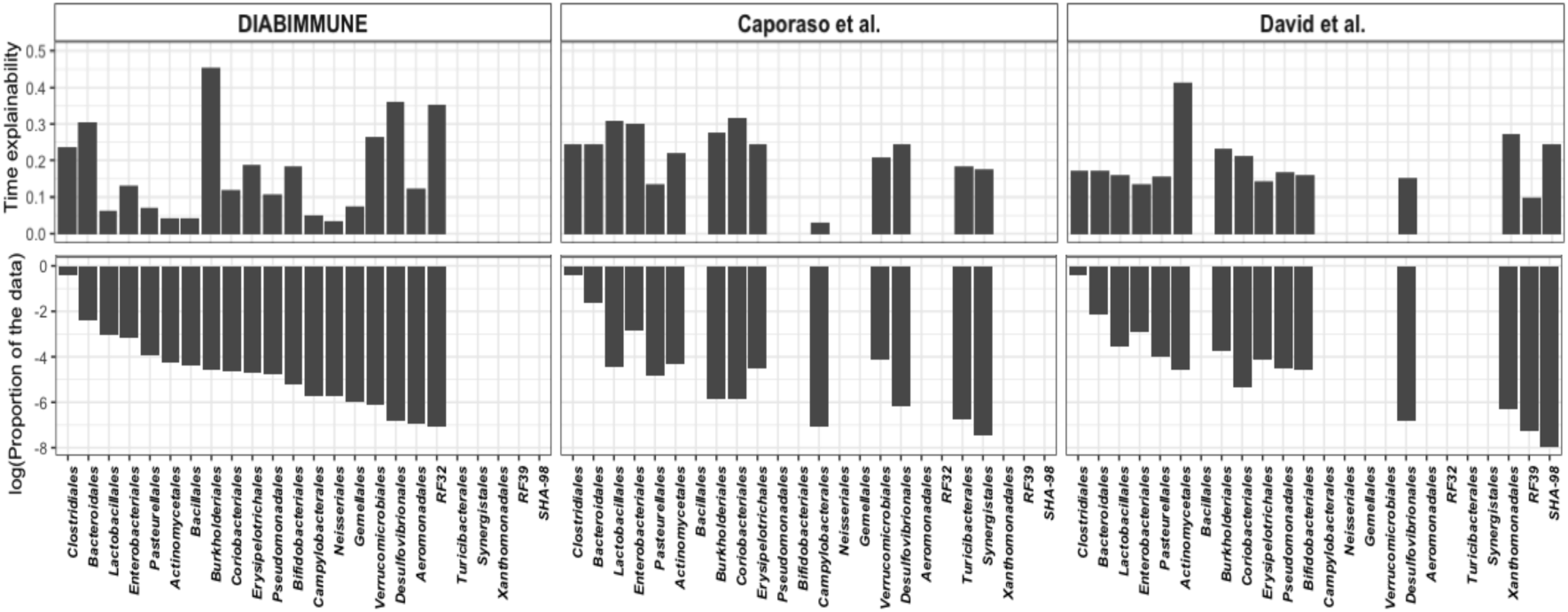
Differential time-explainability of OTUs aggregated by the taxonomic order across all datasets. In the top row, the y-axis is the average time-explainability and in the bottom row, the y-axis is the proportion of OTUs per order in log scale. The x-axis shows orders with OTUs that are autoregressive across all datasets.

Particularly, in the DIABIMMUNE infant data set, there are 7244 OTUs which are divided into 55 different orders. However, the OTUs recognized by *MTV-LMM* as autoregressive (1387 out of 7244) are represented in only 19 orders out of the 55. The remaining 36 orders do not include any autoregressive OTUs. Unlike the autoregressive dynamics, these non-autoregressive dynamics carry a strong phylogenetic structure (P-value *<* 10^*−*16^), that may indicate a niche/habitat filtering. This observation is consistent with the findings of Gibbons et al. [24], who found a strong phylogenetic structure in the non-autoregressive dynamics in the adult microbiome. We validated this finding using the adult microbiome datasets (Supplementary).

Notably, across all datasets, there is no significant correlation between the order dominance (number of OTUs in the order) and the magnitude of its time-explainability component (median *r* = 0.12). For example, in the DIABIMMUNE data set, the proportion of autoregressive OTUs within the 19 mixed orders varies between 2% and 75%, where the average is approximately 20%. In the most dominant order, Clostridiales (representing 68% of the OTUs), approximately 20% of the OTUs are autoregressive and the average time-explainability is 23%. In the second most dominant order, Bacteroidales, approximately 35% of the OTUs are autoregressive and the average time-explainability is 31%. In the Bifidobacteriales order, approximately 75% of the OTUs are autoregressive, and the average time-explainability is 19% (Fig. 4). We hypothesize that the high percentage of autoregressive OTUs in the Bifidobacteriales order, specifically in the infants dataset, can be at least partially attributed to the finding made by [35], according to which some sub-species in this order appear to be specialized in the fermentation of human milk oligosaccharides and thus can be detected in infants but not in adults. This emphasizes the ability of *MTV-LMM* to identify taxa that have prominent temporal dynamics that are both habitat and host-specific.

As an example of *MTV-LMM*’s ability to differentiate autoregressive from non-autoregressive OTUs within the same order, we examined Burkholderiales, a relatively rare order (less than 2% of the OTUs in the data) with 76 OTUs overall, where only 19 of which were recognized as auto-regressive by *MTV-LMM*. Indeed, by examining the temporal behavior of each non-autoregressive OTU in this order, we witnessed abrupt changes in the OTU abundance over time, where the maximal number of consecutive time points with abundance greater than 0 is very small. On the other hand, in the autoregressive OTUs, we witnessed a consistent temporal behavior, where the maximal number of consecutive time points with abundance greater than 0 is well over 10 (Supplementary Fig. S5).

Next, we analyzed the adult microbiome datasets (Caporaso et al. and David et al.) and observed that the temporal presence-absence patterns of the OTUs, when aggregated into taxonomic families, are highly associated with the autoregressive component. Specifically, we found a significant linear correlation between the time-explainability and the family time-prevalence, defined as the median number of time points in which the family OTUs are present (*r* = 0.74, 0.41 respectively).

This correlation was not as strong in the DIABIMMUNE infant dataset (*r* = 0.19), which exhibits a higher variance in time-prevalence within families, a finding that is consistent with known differences between the relatively stable adult microbiome and the developing infant microbiome. Notably, the autoregressive OTUs, across datasets, are significantly more prevalent over time in comparison to the non autoregressive OTUs (P-value *<* 10^*−*16^).

### Communities with high temporal changes in alpha-diversity are prone to a higher autoregressive component

In our previous analysis, we found that the fraction of autoregressive OTUs varies across different datasets. In order to shed light on the underlying factors that may lead to temporal dependencies, we compared the autoregressive components across the different datasets by examining the number of autoregressive OTUs, the time-explainability of each OTU and the correlation with the measured abundance. We find a positive correlation between the time-explainability and the first derivative of the alpha-diversity over time. Specifically, communities with high temporal changes in alpha-diversity are prone to a larger autoregressive component. These findings are supported by [28] which states that microbial communities undergoing frequent changes, such as that of infants, are the most informative for temporal dynamic modeling. On the other hand, adult microbial communities often display stability and resilience, which leads to measurements mostly consistent with a steady-state.

Indeed, *MTV-LMM*’s estimation of the time-explainability component (23%) is higher in the DIABIMMUNE dataset in comparison to the adult datasets. The DIABIMMUNE dataset is fundamentally different from the other datasets since it measures the development of the gut microbial community in infants, and is characterized by significant temporal changes in alpha-diversity. On the other hand, the David et al. data presented with a relatively stable alpha-diversity over time, and with an average estimated time-explainability of 14% (Fig. 5). Notably, despite the relatively stable nature of the adult microbiome, *MTV-LMM* can nonetheless identify autoregressive OTUs and therefore presented with improved prediction accuracy compared to other methods (Fig. 1).

**Figure 5.**
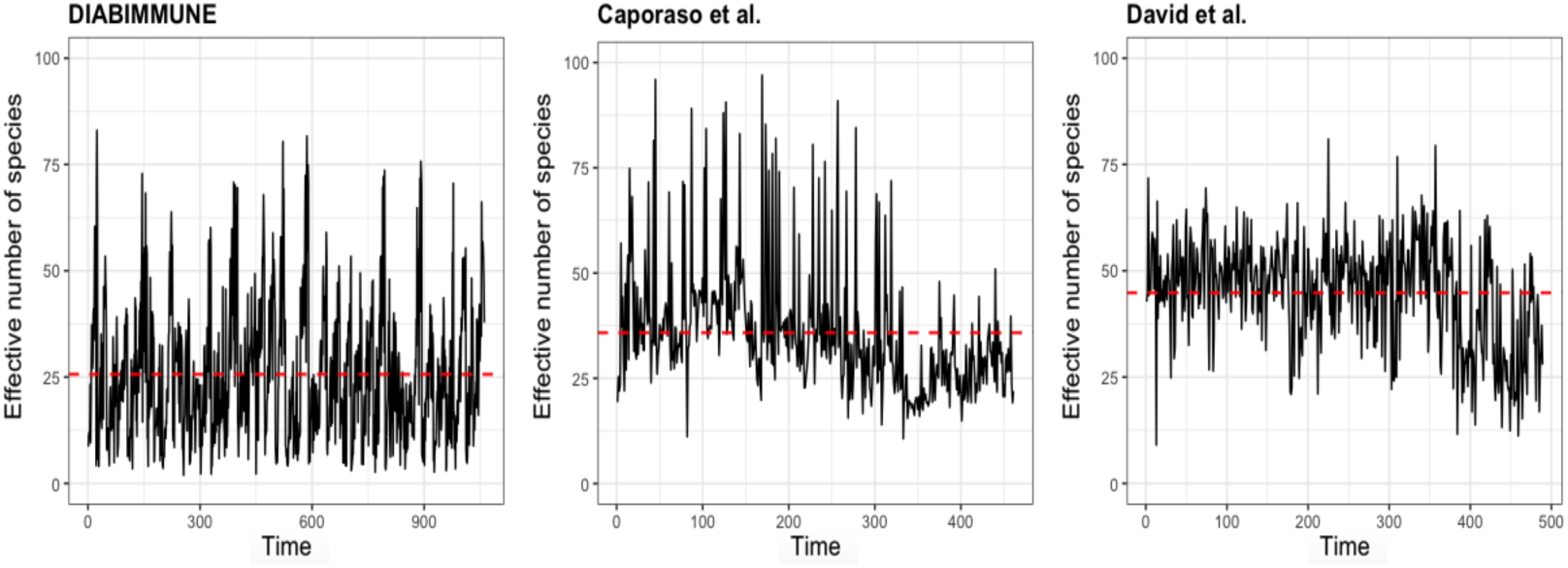
Black lines show the Shannon ‘effective number of species’ *N*_*eff*_ (a measure of alpha diversity) for each dataset. The red dashed lines show the average *N*_*eff*_ for each dataset.

### The autoregressive component of an adult versus an infant microbiome

The colonization of the human gut begins at birth and is characterized by a succession of microbial consortia [36–39], where the diversity and richness of the microbiota reach adult levels in early childhood. A longitudinal observational study has recently been used to show that infant gut microbiota begins transitioning towards an adult-like community after weaning [40], implying that one of the most dominant factors associated with variation in an infant microbiome is time. This observation is validated using our infant longitudinal data set (DIABIMMUNE) by applying PCA on the temporal kinship matrix (Fig. 6). Our analysis reveals that the first principal component (accounting for 26% of the overall variability) is associated with time. Specifically, there is a clear clustering of the time samples from the first six months of an infant’s life and the rest of the time samples (months 7 *−* 36) which can be correlated to weaning. As expected, we find a strong autoregressive component in an infant microbiome, which is highly associated with temporal variation across individuals. Using *MTV-LMM*, we utilized the high similarity between infants over time, and simultaneously quantify the autoregressive component in their microbiome.

**Figure 6.**
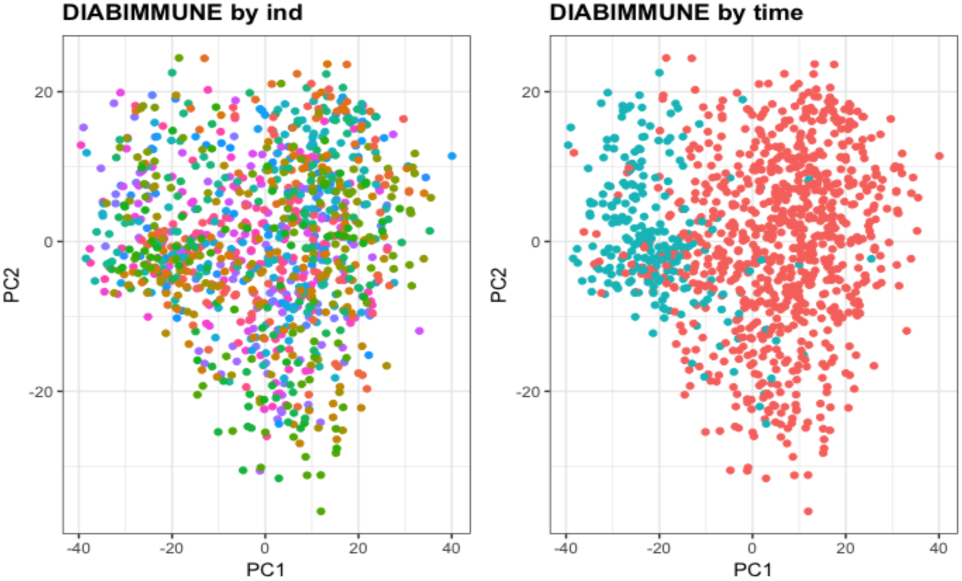
The first two PCs of the temporal kinship matrix color coded by individual (left) and by time - before and after six months (right) in the DIABIMMUNE data (39 infant donors).

In contrast to the infant microbiome, the adult microbiome is considered relatively stable [16, 41], but with considerable variation in the constituents of the microbial community between individuals. Specifically, it was previously suggested that each individual adult has a unique gut microbial signature [42–44], which is affected, among others factors, by environmental factors [20] and host lifestyle (i.e., antibiotics consumption, high-fat diets [17] etc.). In addition, [17] showed that over the course of one year, differences between individuals were much larger than variation within individuals. This observation was validated in our adult datasets (David et al. and Caporaso et al.) using the same PCA of the temporal kinship matrix which reveals that the first principal component, which accounts for 61% and 43% of the overall variability respectively, is associated with the individual’s identity (Fig. 7, Supplementary Fig. 6).

**Figure 7.**
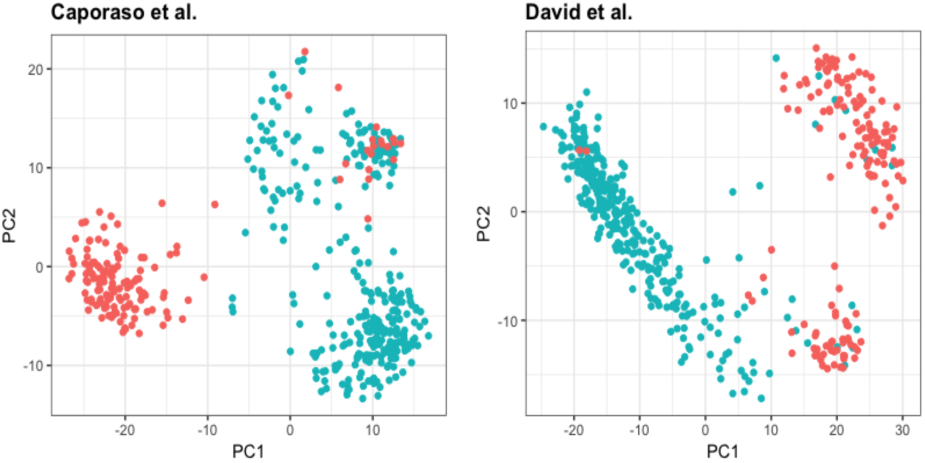
The first two PCs of the temporal kinship matrix color coded by individual in the adults datasets (a) Caporaso et al. (2 adult donors) (b) David et al. (2 adult donors).

Using *MTV-LMM* we observe that despite the large similarity along time within adult individuals, there is also a non-negligible autoregressive component in the adult microbiome. The fraction of variance explained by time across individuals can range from 6% up to 79%. These results shed more light on the temporal behavior of species in the adult microbiome as opposed to that in infants, which are known to be highly affected by time [40]. To the best of our knowledge, there is no prior evidence of a significant autoregressive component in adult microbiomes.

## Methods

### The *MTV-LMM* Algorithm

*MTV-LMM* uses a linear mixed model (see [45] for a detailed review), a natural extension of standard linear regression, for the prediction of time series data. We describe the technical details of the linear mixed model below.

We assume that the relative abundance levels of a specific OTU *j* at time point *t* depend on a linear combination of the relative abundance levels of the OTUs in the microbial community at previous time points. We further assume that the temporal changes in relative abundance levels, of a specific OTU *j*, are a time-homogeneous high-order Markov process. We model the transitions of this Markov process using a linear mixed model, where we fit the *p* previous time points of OTU *j* as fixed effects and the *q* previous time points of all other OTUs as random effects. *p* and *q* are the temporal parameters of the model.

For simplicity of exposition, we present the generative linear mixed model that motivates the approach taken in *MTV-LMM* in two steps. In the first step we model the time series for each OTU in one individual host. In the second step we extend our model to *N* individuals, while accounting for the hosts’ effect.

We first describe the model assuming there is only one individual. We assume that the relative-abundance levels of *m* OTUs, denoted as the microbial community, have been measured at *T* time points. We get as input an *m × T* matrix *M*, where *M_jt_* represents the relative-abundance levels of OTU *j* at time point *t*. Let *y*^*j*^ = (*M*_*j,p*+1_*,…, M*_*jT*_)^*t*^ be a (*T − p*) *×* 1 vector of OTU *j* relative abundance, across *T − p* time points starting at time point *p* + 1 and ending at time point *T*. Let *X*^*j*^ be a (*T − p*)*×*(*p* + 1) matrix of *p* covariates, comprised of an intercept vector as well as the first *p* time lags of OTU *j* (i.e., the relative abundance of OTU *j* in the *p* time stamps prior to the one predicted). Formally, for *k* = 1 we have 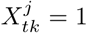, and for *p* + 1 *≥ k >* 1 we have 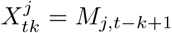. For simplicity of exposition and to minimize the notation complexity, we assume for now that *p* = 1. Let *W* be an (*T − q*) *× q ⋅ m* standardized relative abundance matrix, representing the first *q* time lags of the microbial community. For simplicity of exposition we describe the model in the case *q* = 1, and then *W*_*tj*_ = *M*_*jt*_ (in the more general case, we have *W*_*tj*_ = *M*_⌈*j/q⌉,t*−(*j mod q*)_).

With these notations, we assume the following linear model:

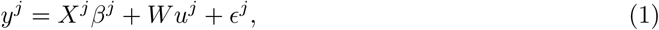

where *u*^*j*^ and *∈*^*j*^ are independent random variables distributed as 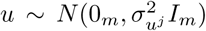 and 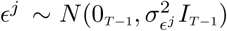. The parameters of the model are *β*^*j*^ (fixed effects), 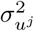, and 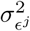.

We note that environmental factors known to be correlated with OTU relative abundance levels (i.e., diet, antibiotic usage [17, 20]) can be added to the model as fixed linear effects (i.e., added to the matrix *X*^*j*^).

Since the relative abundance levels are highly variable across different OTUs, this model is not going to perform well in practice. Intuitively, we would like to capture the information as to whether an OTU is present or absent, or potentially introduce a few levels (i.e., high abundance, medium, and low). We use the quantiles of each OTU to transform the matrix *M* into a matrix 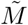, where 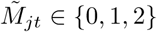 depending on whether the abundance level is low (below 25% quantile), medium, or high (above 75% quantile). We also tried other normalization strategies, including quantile normalization, which is typically used in gene expression eQTL analysis [46, 47], and the results were qualitatively similar (see Supplementary Fig. S6). We subsequently replace the matrix *W* by a matrix 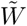, which is constructed analogously to *W*, but using 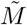 instead of *M*. Thus, our model can now be described as

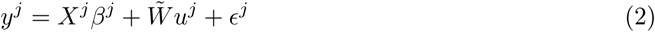

So far, we described the model assuming we have time series data from one individual. We next extend the model to the case where time series data is available from multiple individuals. In this case, we assume that the relative abundance levels of *m* OTUs, denoted as the microbial community, have been measured at *T* time points across *N* individuals. We assume the input consists of *N* matrices, *M*^1^*,…,M*^*N*^, where matrix *M*_*i*_ corresponds to individual *i*, and it is of size *m × T*. Therefore, the outcome vector *y*^*j*^ is now an *n ×* 1 vector. For example, when we describe the dimensions of *∈*_*j*_, composed of *N* blocks, where *n* = (*T −* 1)*N*, where block *i* corresponds to the time points of individual *i*. Formally, 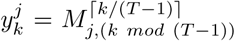. Similarly, we define *X*^*j*^ and 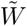 as block matrices, with *N* different blocks, where each block corresponds to individual *i*.

When applied to multiple individuals, Model (2) may overfit to the individual effects, e.g., due to the host genetics. In other words, since our goal is to model the changes in time, we need to condition these changes in time on the individual effects, that are unwanted confounders for our purposes. We therefore construct a matrix *H* by randomly permuting the rows of each block matrix *i* in 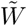, where the permutation is conducted only within the same individual. Formally, we apply permutation *π_i_ ∈ S*_*T−1*_ on the rows of each block matrix *i*, *M*^*i*^, corresponding to individual *i*, where *S*_*T−1*_ is the set of all permutations of (*T −* 1) elements. In each *π*_*i*_, we are simultaneously permuting the entire microbial community. Hence, matrix *H* corresponds to the data of each one of the individuals, but with no information about the time (since the data was shuffled across the different time stamps). With this addition, our final model is given by

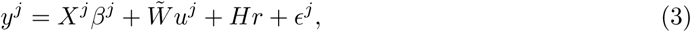

where 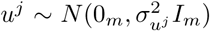 and 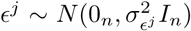, and 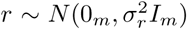. It is easy to verify that an equivalent mathematical representation of model 3 can be given by

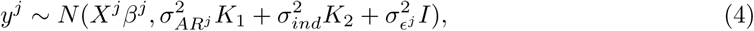

where 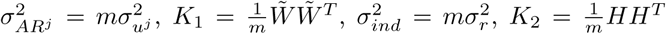. We will refer to *K*_1_ as the *temporal kinship matrix*, or the temporal relationship matrix.

We note that for the simplicity of exposition, we assumed so far that each sample has the same number of time points *T*, however in practice the number of samples may vary between the different individuals. It is easy to extend the above model to the case where individual *i* has *T*_*i*_ time points, however the notations become cumbersome; the implementation of *MTV-LMM*, however takes into account a variable number of time points across the different individuals.

Once the distribution of *y*^*j*^ is specified, one can proceed to estimate the fixed effects *β*^*j*^ and the variance of the random effects using maximum likelihood approaches. Specifically, the common approach for estimating variance components is known as restricted maximum likelihood (REML). We followed the procedure described in the GCTA software package [48], originally developed for genotype data, and adjusted it for the OTU data. GCTA implements the REML method via the average information (AI) algorithm.

### Time-explainability

We define the term *time-explainability*, denoted as *χ*, to be the variance of a specific OTU relative abundance explained by the microbial community in the previous time points. Formally, for OTU *j* we define

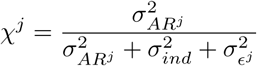

The time-explainability was estimated with GCTA, using the temporal kinship matrix. Confidence intervals were computed using FIESTA [49]. In order to measure the accuracy of time-explainability estimation, the average confidence interval width was estimated by computing the confidence interval widths for all auto-regressive OTUs and averaging the results. Additionally, we adjust the time-explainability P-values for multiple comparisons using the Benjamini-Hochberg method [50].

### Best Linear Unbiased Predictor

We now turn to the task of predicting 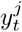 using the OTU relative abundance in time *t −* 1 (or more generally in the last few time points). Using our model notation, we are given *x*^*j*^ and 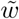, the covariates associated with a newly observed time point *t* in OTU *j*, and we would like to predict 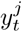 with the greatest possible accuracy. For a simple linear regression model, the answer is simply taking the covariate vector *x* and multiplying it by the estimated coefficients 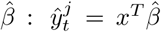. This practice yields unbiased estimates. However, when attempting prediction in the linear mixed model case, things are not so simple. One could adopt the same approach, but since the effects of the random components are not directly estimated, the vector of covariates 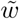 will not contribute directly to the predicted value of 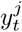, and will only affect the variance of the prediction, resulting in an unbiased but inefficient estimate. Instead, one can use the correlation between the realized values of 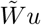, to attempt a better guess at the realization of 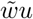 for the new sample. This is achieved by computing the distribution of the outcome of the new sample conditional on the full dataset, by using the following property of the multivariate normal distribution. Assume we sampled *t−* 1 time points from OTU *j*, but the relative abundance level for next time point *t*, 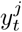, is unknown. The conditional distribution of 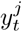 given the relative abundance levels at all previous time points, *y*^*j*^, is given by:

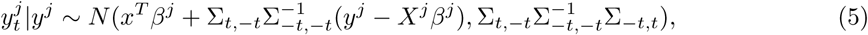

where 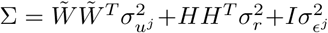 and positive/negative indices indicate the extraction/removal of rows or columns, respectively. Intuitively, we use information from the previous time points that have a high correlation with the new time point, to improve its prediction accuracy. The practice of using the conditional distribution is known as BLUP (Best Linear Unbiased Predictor). Therefore, *MTV-LMM* could be used to learn OTU effects in a discovery set (OTU table at time points 1,…,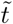), and subsequently use these learned OTU effects to predict the temporal-community contribution in the next time point (OTU *j* at 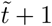). In our experiments, the initial 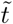 was set to be approximately 1*/*3 of the time series length in each data set.

### Prediction accuracy

The predictive ability of a model is commonly assessed using the prediction error variance, 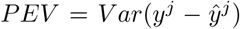. The proportional reduction in relative abundance variance accounted for by the predictions (referred to as *R*^2^ in this paper) can be quantified using

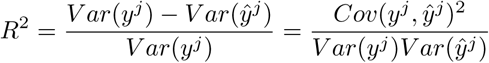

### Model selection

Given that the model presented in equation (3) can be extended to any arbitrary *p* and *q*, we tested four different variations of this model: 1. *p* = 0 and *q* = 1 (no fixed effect, one random effect based on 1-time lag), 2. *p* = 1 and *q* = 1 (one fixed effect based on 1-time lag, one random effect based on 1-time lag), 3. *p* = 0 and *q* = 3 (no fixed effect, one random effect based on 3-time lags) and 4. *p* = 1 and *q* = 3 (one fixed effect based on 1-time lag, one random effect based on 3-time lags).

We divide each dataset into three parts - training, validation, and test, where each part is approximately 1*/*3 of the time series (sequentially). We train all four models presented above and use the validation set to select a model for each OTU *j* based on the highest correlation with the real relative abundance. We then compute sequential out-of-sample predictions on the test set with the selected model.

There are three main justifications for the use of multiple time points in the model. First, Gibbons et al. [24] empirically preformed a time-lag analysis and found that according to the autocorrelation decay curves, most of the autocorrelation was gone after a lag of 3 or 4 days, whereas some of the autocorrelation was gone from the time series after 1 or 2 days. Second, several peer-reviewed articles [51–54] found that the human microbiome reaches equilibrium within 10 days following small perturbations to the community. It is imperative to model the different OTUs in a manner that will fit their temporal patterns. Third, allowing for the use of multiple previous timestamps increases flexibility so that the model can select the correct time window required for each OTU.

### Phylogenetic analysis

We performed the following phylogenetic analysis. First, in order to test the hypothesis that both auto-regressive and non-autoregressive dynamics carry a taxonomic signal, we fitted a linear mixed model, where the kinship matrix is the phylogenetic distance between pairs of OTUs and the outcomes are the time-explainability measurement for each OTU. Second, in order to test the hypothesis that only non-autoregressive dynamics carry a taxonomic signal that is better than random, we conducted a permutation test by shuffling the taxonomic order assigned to each OTU - generating new random “orders” - over 100, 000 iterations. We counted the number of non-autoregressive orders in each iteration, thereby generating a sampling distribution, which we then used to calculate an exact P-value for the dataset in each iteration. Lastly, we calculated the linear correlation between ‘time-prevalence’ - defined as the median number of time points in which the OTU is present - and time-explainability for each taxonomic family.

### Alpha diversity measures

To measure the alpha diversity, we used Shannon-Wiener index, which is defined as *H* = − Σ*p_j_ln*(*p*_*j*_), where *p*_*j*_ is the proportional abundance of species *j*. Shannon-Wiener index accounts for both abundance and evenness of the species present. Additionally, we computed the ‘effective number of species’ (also known as true diversity), the number of equally-common species required to give a particular value of an index. The ‘effective number of species’ associated with a specific Shannon-Wiener index ‘*a*’ is equal to *exp*(*a*).

### Preliminary OTU screening according to temporal presence-absence patterns

For calculating the temporal kinship matrix we included OTUs using the following criteria. An OTU was included if it was present in at least 0.1% of the time points (removes dominant zero abundance OTUs). In the David et al. dataset we included 1051 (out of 2804), in the Caporaso et al. dataset we included 922 (out of 3436) and in the DIABIMMUNE dataset we included 1440 (out of 7244) OTUs.

### Methods comparison

We compared *MTV-LMM* to two existing methods: sVAR suggested by [24] and Poisson regression suggested by [29]. In the sVAR method, we followed the procedure described in [24], while running the model and computing the prediction for each individual separately. We then computed an aggregated prediction accuracy score for each OTU, by averaging the prediction accuracy of each individual, using the OTUs the passed the preliminary screening as described above. In the Poisson regression method, we followed the procedure described in [29], while running the model for all the individuals simultaneously and calculating prediction accuracy for each OTU. We used the OTUs that passed the screening suggested in [29] (eliminating any OTUs in the data for which there were a small number (*<* 6) of average reads per sample). In both models, the training set was 0.67 of the data and the test set was the remaining 0.33 of the data. In both cases we used the code supplied by the authors.

### Datasets

We evaluated the performance of *MTV-LMM* using three datasets collected using 16S rRNA gene sequencing. All data sets are publicly available. The first data set was collected and studied by David et al. (2014) [17] (2 adult donors). The next data set was collected and studied by Caporaso et al. (2011) [16] (2 adult donors). The third data set was collected by the ‘DIABIMMUNE’ project and studied by Yassour et al. (2016) [21] (39 infant donors). In order to compare across studies and reduce technical variance between studies, closed reference OTUs were clustered at 99% identity against the Greengenes database *v.*13_5_*/*8 [55]. Open reference OTU picking was also run [56], in order to look for non-database OTUs that might contribute substantially to community dynamics. Time series OTU tables were normalized by random sub-sampling to contain 10, 000 reads per sample.

*David et al. (2014)* dataset [17]. Stool samples from 2 healthy American adults were collected (donor A = DA and donor B = DB). DA collected gut microbiota samples between days 0 and 364 of the study (total 311 samples). DB primarily collected gut microbiota samples between study days 0 and 252 (total 180 samples). The V4 region of the 16S ribosomal RNA gene subunit was used to identify bacteria in a culture-independent manner. DNA was amplified using custom barcoded primers and sequenced with paired-end 100 bp reads on an Illumina GAIIx according to a previously published protocol [57]. ‘OTU picking’ and ‘quality control’ were performed essentially as described [17]. In this work, we used the OTUs shared across donors (2, 804 OTUs).

*Caporaso et al. (2011)* dataset [16]. Two healthy American adults, one male (M3) and one female (F4), were sampled daily at three body sites (gut (feces), mouth, and skin (left and right palms)). M3 was sampled for 15 months (total 332 samples) and F4 for 6 months (total 131 samples). Variable region 4 (V4) of 16S rRNA genes present in each community sample were amplified by PCR and subjected to multiplex sequencing on an Illumina Genome Analyzer IIx according to a previously published protocol [57]. ‘OTU picking’ and ‘quality control’ were performed essentially as described [16]. In this work, we used the OTUs shared across donors (3, 436 OTUs).

*DIABIMMUNE* dataset [21]. Monthly stool samples from 39 Finnish children aged 2 to 36 months (total of 1101 samples). To analyze the composition of the microbial communities in this cohort, DNA from stool samples was isolated and amplified and sequenced the V4 region of the 16S rRNA gene. Sequences were sorted into OTUs. 16S rRNA gene sequencing was performed essentially as previously described in [21]. In this work, we used all the OTUs in the sample (7, 244 OTUs).

## Discussion

We have presented *MTV-LMM*, a method for modeling and analyzing time series microbial community data. *MTV-LMM* can be used to predict the abundance levels of OTUs in the future, and it can also be used to assess the time-explainability of each of the OTUs, i.e., how much of the variability of the OTU abundance levels can be explained based on past microbial community composition. Time-explainability can be informative for selecting autoregressive OTUs that are essential to understanding microbiome behavior in longitudinal studies. In particular, such OTUs can be used to characterize temporal trajectories of the microbial community. This provides a stepping stone towards a better understanding of microbial community temporal dynamics as well as the use of microbiome data as a diagnostic or forensic tool in various fields like healthcare or epidemiology.

We have demonstrated that *MTV-LMM* significantly outperforms existing approaches for temporal modeling of the microbiome, both in terms of its prediction accuracy’as well as in its ability to identify time-dependent OTUs. We hypothesize that this improved performance stems from the fact that unlike previous approaches, *MTV-LMM* leverages the information across an entire population, while adjusting for the individual’s effect, i.e. genetic and environmental effects that affect each individual differently.

*MTV-LMM* is a flexible and computationally efficient tool, which can be easily adapted by researchers to select the core time-dependent OTUs, quantify their temporal effects (given the microbial community composition in previous time points) and predict their future abundance. Additionally, as we demonstrate in the Results section (Fig. 2), the interactions estimated by *MTV-LMM* can be used to uncover global patterns in the microbial composition dynamics. Using the DIABIMMUNE dataset, we found a strong phylogenetic structure in the species-species interaction matrix estimated by *MTV-LMM*.

Using *MTV-LMM*, we have demonstrated that OTU autoregressiveness is a spectrum where certain OTUs are almost entirely determined by the community composition at previous time points, some are somewhat dependent on the previous time points, and others are completely independent of previous time points. We further show that this autoregressive characterization is related to alpha diversity and to the phylogenetic structure of the microbial community. The phylogenetic orders harboring autoregressive OTUs are of a mixed nature, containing both autoregressive and non-autoregressive OTUs in different proportions (Fig. 4). Notably, the ability of *MTV-LMM* to differentiate between OTUs within the same order can be utilized to find keystone species that may be responsible for the temporal changes observed in the different datasets.

By applying *MTV-LMM* to three temporal datasets from the gut microbiome [16,17,21], we found that, unlike previously thought, a considerable proportion of OTUs have a non-negligible autoregressive component in both infants and adults. Specifically, we show that on average, the autoregressiveness measured by the time-explainability is an order of magnitude larger than previously appreciated. Our results provide such evidence of autoregressiveness in the adult microbiome for the first time, a finding which was missed by previous approaches due to underestimation of the time-explainability of OTUs. This highlights the sensitivity of our method and its ability to handle both developing and stable datasets. Furthermore, our method and findings suggest that a considerable proportion of the human gut microbiome’s temporal behavior can be predicted with high accuracy. These results are an important step towards better prediction and possibly better manipulation of the microbiome composition in adults in the future.

As shown in the Results section (Fig. 6), applying PCA on the temporal kinship matrix in the infant dataset revealed a clear clustering of the time samples that separated the first six months of an infant’s life from the rest of the time samples (7-36 months). We hypothesize that a major dietary change was the cause for this temporal clustering – most likely weaning. Indeed, our results correspond to a recent longitudinal observational study showing that the infant gut microbiota begin transitioning towards adult communities after weaning, roughly at the age of six months [40]. Furthermore, our results are in agreement with those presented by [22] in relation to dietary effects, which validate the ability of *MTV-LMM* to detect the relevant OTUs affected by dietary changes in the developing microbiome.

It is important to note that the time-explainability of a given OTU may strongly depend on the density of the sampling. For example, if the microbiome is sampled every two months as opposed to every month, the time-explainability may be reduced since more noise is added over time (i.e., fewer environmental changes are tracked). The instrumental novelty of our method to predict a specific OTU’s temporal behavior is the statistical power that is gained by leveraging the overall community composition as well as all the individuals in the dataset. This suggests that interactions within the microbiome are of major importance in modulating a specific microbe’s behavior over time.

